# Network Modulation Enables 3D-Printed Citrate-Based Polymer Scaffolds with Broadly Tunable Mechanical Performance for Regenerative Engineering

**DOI:** 10.64898/2026.02.12.705586

**Authors:** Ni Chen, Nolan Schlessman, Rao Fu, Yonghui Ding

## Abstract

Biomaterials with highly tunable mechanical properties and tissue-mimetic structural features are critical for diverse biomedical applications. Photopolymerizable citrate-based polymers (CBP), such as methacrylate polydiolcitrate (mPDC), enable high-resolution fabrication of biodegradable scaffolds via light-based 3D printing for regenerative engineering. However, mPDC scaffolds typically exhibits substantial brittleness due to the formation of highly crosslinked and heterogeneous polymer network, an intrinsic limitation of many acrylate-based polymers, thereby restricting their use across a broad range of tissue types. Herein, we report facile network-engineering strategies to modulate crosslinking density and network topology of CBPs through the incorporation of acrylate-based reactive diluents and/or a thiol-based chain transfer agent, 3,6-dioxa-1,8-octanedithiol (DOD). These approaches enabled significantly improved and broadly tunable mechanical properties, with Young’s modulus spanning 6.8-134 MPa, ultimate tensile strength ranging from 1.8 MPa to 18 MPa, and strain at break varying from 14% to 61%. Notably, incorporation of isobornyl acrylate (IBOA) alone significantly enhanced toughness, yielding a 3.6-fold increase in Young’s modulus (50 MPa vs. 14 MPa) and a 2.8-fold increase in strain at break (39% vs. 14%). Moreover, combined incorporation of IBOA and DOD remarkably improved ductility, achieving a 4-fold increase in strain at break to 61% while maintaining comparable stiffness. All mPDC composites exhibited tunable biodegradability, good cytocompatibility, and excellent 3D printability. Using these composite inks, 3D-printed meniscus scaffolds supported the human chondrocyte growth and fibrochondrogenic matrix deposition, while 3D-printed vascular stents supported endothelial monolayer formation. Collectively, this study establishes a versatile photopolymerizable citrate-based biomaterial platform with broadly tunable mechanical performance, controllable biodegradability, good cytocompatibility, and high printability, offering strong potential for customized biomedical applications ranging from load-bearing to soft tissue engineering.

## 1. Introduction

Regenerative engineering often relies on biomaterial scaffolds to repair or restore mechanical and biological functions of damaged tissues and organs. To achieve functional integration, scaffolds must closely recapitulate the mechanical and structural characteristics of native tissues, thereby ensuring mechanical stability, structural integrity, and effective tissue integration.^1, 2^ Importantly, the mechanical requirements of scaffolds vary substantially across tissue types. For example, scaffolds designed for musculoskeletal tissue repair, such as tendon and ligament, require high stiffness with moduli reaching to several hundred of MPa.^3–7^ In contrast, scaffolds for soft tissue regeneration, such as blood vessels, must exhibit high flexibility, typically with moduli of 1-2 MPa and strain at break ranging from 24% to 150 %.^8, 9^ In addition to mechanical performance, scaffold geometry and porous structure play critical roles in regulating scaffold-tissue integration. Three-dimensional (3D) printing provides a powerful approach to fabricate scaffolds with anatomically accurate geometry and well-defined porous structures, enabling improved tissue integration and favorable biological responses.^10, 11^ Among various 3D printing techniques, photopolymerization-based 3D printing methods, such as continuous liquid interface production (CLIP), offer an favorable combination of printing speed, resolution, and cost efficiency.^12–14^ However, photopolymerization-based 3D printing often fails to deliver scaffolds with broadly tunable mechanical properties suitable for diverse target tissues, largely due to the limited availability of photopolymerizable biomaterial inks.

Synthetic biodegradable polymer materials have been widely used for scaffold fabrication due to their controllable mechanical properties and degradation behavior, as well as their compatibility with diverse manufacturing approaches. To date, ten biodegradable polymers have been used in implantable medical devices cleared or approved by U.S. Food and Drug Administration (FDA). Nevertheless, many of these polymers lack intrinsic bioactivity to support cell adhesion, differentiation and tissue regeneration.^15^ Moreover, degradation products of poly (glycolic acid) (PGA) and poly (lactic acid) (PLA)-based polymers, i.e. glycolic acid and lactic acid, can accumulate at the implantation sites and trigger chronic inflammation.^16^

In recent years, citrate-based polymers (CBPs), such as polydiolcitrates, have attracted great attention and have been commercially developed due to their unique combination of biocompatibility, biodegradability, antioxidant/anti-inflammatory properties.^17–19^ However, conventional CBPs are typically cured through heat-induced polycondensation, a process that often requires days of reaction time in molds, limiting fabrication efficiency and restricting manufacturability for scaffolds with complex geometries. To overcome this limitation, photopolymerizable polydiolcitrates have been developed through methacrylation, yielding methacrylate polydiolcitrates (mPDC) that enable rapid fabrication of complex scaffolds via CLIP-based 3D printing.^20^ These scaffolds demonstrated high stiffness, excellent biocompatibility, mild inflammatory responses, and minimal restenosis following implantation in swine coronary arteries over four weeks.^20^ Despite these advantages, photopolymerized mPDC exhibits pronounced brittleness, with elongation at break typically below 15%, markedly lower than that of their polycondensed counterparts.^21^ This brittleness represents an intrinsic limitation of many acrylate-based polymers and is attributed to high crosslinking density and network heterogeneity resulting from rapid photopolymerization kinetics.^22^ A recent study incorporated highly elastic natural rubber into a polycondensed CBP network, which increased strain at break from 88% to 415%.^23^ However, this ductility enhancement was not retained in photopolymerized, methacrylate CBP systems, where strain at break decreased to only 17-20%. Together, these findings underscore a persistent challenge: while photopolymerizable acrylate-based CBP biomaterials enable rapid fabrication of complex scaffolds through 3D printing, their inherent brittleness remains a major barrier to broad applications in regenerative engineering.

To address this challenge, we report two facile network-engineering strategies to effectively modulate crosslinking density and network topology in photopolymerizable CBP inks by incorporating mono-functional acrylate-based reactive diluents (RDs) and/or a thiol-based chain transfer agent into the methacrylate CBP formulations. These strategies enable significantly improved and highly tunable mechanical performance while preserving rapid and robust photopolymerization-based 3D printability. The physiochemical properties, mechanical performance, degradation behavior, and cytocompatibility of photopolymerized CBPs containing RDs and/or thiol molecules were systematically characterized to elucidate structure-property relationships. Furthermore, we demonstrated the successful fabrication of meniscus scaffolds and vascular stents using CBP inks and CLIP-based 3D printing, as well as their ability to support fibrochondrogenic matrix deposition by human chondrocytes and endothelial monolayer formation, respectively. Collectively, these results highlight the strong potential of this photopolymerizable CBP biomaterial platform for repairing or regenerating a broad-spectrum tissues.

## 2. Results and Discussion

### 2.1 Design of mPDC composite inks with RDs and/or DOD for network modulation

Photopolymerizable CBPs, i.e., methacrylate poly(1,12-dodecanethylene citrate) (mPDC), were synthesized from citric acid and 1,12-dodecanediol following our previously published protocols.^24^ Successful methacrylation was confirmed by ^1^H-NMR spectroscopy, which verified the characteristic chemical shifts associated with both the alkyl backbone and methacrylate double bonds (**Supplementary Information Figure S1**). To modulate the crosslinking density and network topology of photopolymerized mPDC, two network-engineering strategies were employed (**Figure 1**). The first strategy involved incorporating mono-functional acrylate-based RDs, including isobornyl acrylate (IBOA), isodecyl acrylate (IDA), and butyl acrylate (BTA), into the mPDC formulations. These RDs were selected to reduce ink viscosity, which is an important requirement for achieving high-resolution and rapid CLIP-based 3D printing,^22^ and more importantly, to tune the effective crosslinking density of the resulting polymer network (**Figure 1b**). The second strategy introduced a thiol-based chain transfer agent, i.e. 3,6-dioxa-1,8-octanedithiol (DOD), into the mPDC formulations to modify the polymer network topology. Incorporation of DOD was expected to shift the polymerization mechanism towards a mixed-mode that combines methacrylate chain-growth with methacrylate-thiol step-growth polymerization. This mixed-mode polymerization mechanism is anticipated to yield a more homogeneous polymer network and thereby improving the ductility of photopolymerized mPDC (**Figure 1b**).^25, 26^ All composite inks were photopolymerized under UV irradiation (365 nm) or fabricated using a home-built CLIP-based 3D printing system, followed by post-curing at 80°C for 24 h. The resulting polymers were then systematically characterized in terms of physicochemical properties, mechanical performance, degradation behavior, and cytocompatibility.

**Figure 1.**
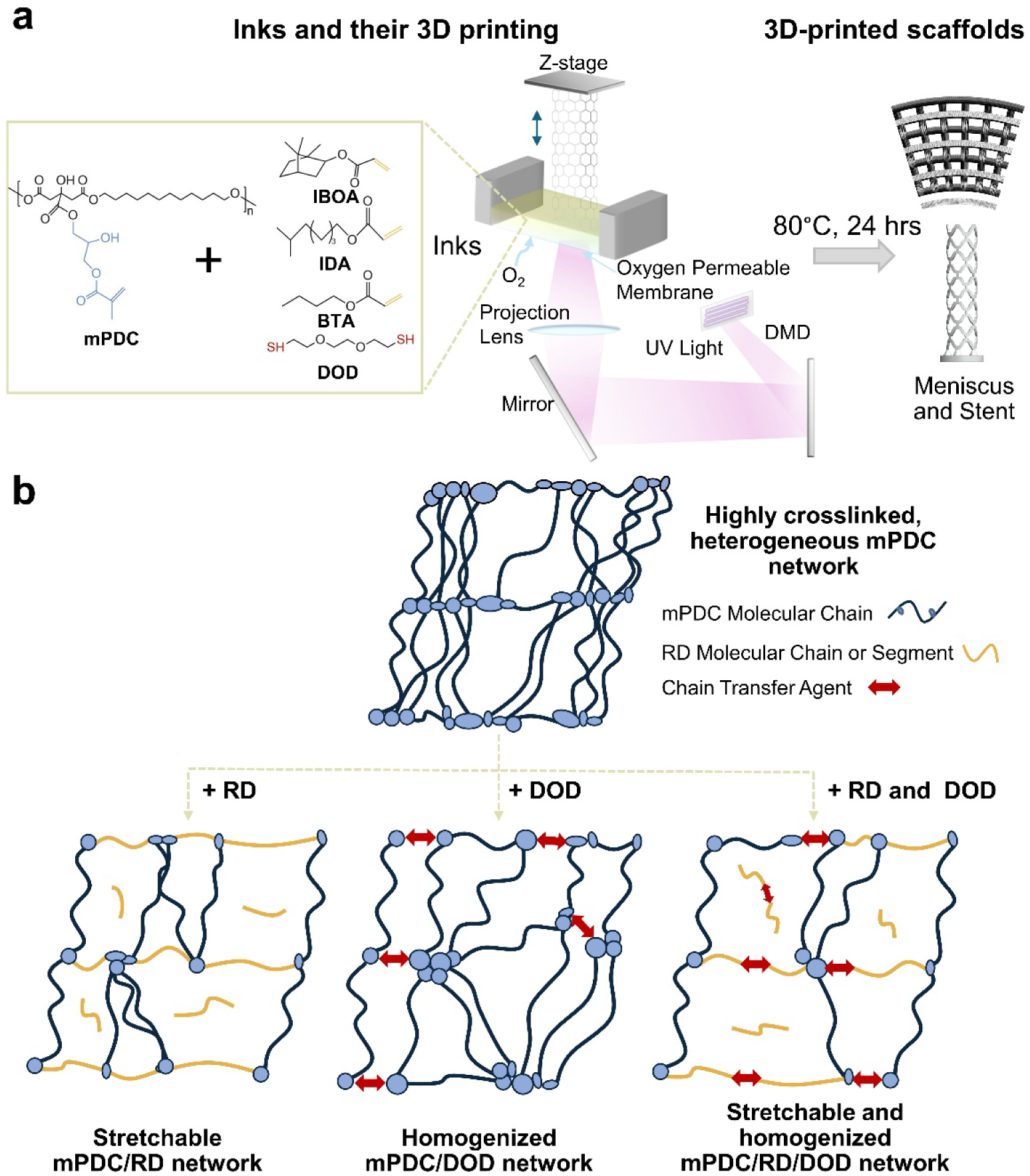
Schematic illustration of network-engineering strategies for modulating photopolymerizable mPDC composite inks through incorporation of RDs and/or a thiol molecule. **(a)** Formulation of mPDC inks containing RDs and/or DOD, followed by CLIP-based photopolymerization 3D printing and post-processing to fabricate 3D scaffolds. **(b)** Transformation of the mPDC network from a highly crosslinked and heterogeneous structure into a stretchable mPDC/RD network, a homogenized mPDC/DOD network, or a stretchable and homogenized mPDC/RD/DOD network through incorporation of RDs and/or DOD.

### 2.2 Chemical structure and thermal properties of mPDC composites

The mPDC/RD composites were prepared by formulating inks containing an equimolar amount of RD relative to the nominal methacrylate content of mPDC, resulting in composite inks containing 20 wt% IBOA, 20 wt% IDA, and 12.5 wt% BTA (**Supplementary Information Table S1**). The chemical structure of photopolymerized mPDC/RD composites were characterized by Fourier-transform infrared spectroscopy (FTIR). For comparison, RD homopolymers were prepared by UV polymerization of individual RD monomers using Irgacure 819 (I819) as the photoinitiator and were similarly analyzed by FTIR. As shown in **Figure 2a**, characteristic adsorption bands of mPDC, including the stretching vibration of hydrogen-bonded hydroxyl groups at 3475 cm^-1, 27^ C-H stretching bonds at 2925 cm^-1^ (symmetric) and 2853 cm^-1^ (asymmetric), ^28^ were clearly observed in all mPDC/RD composites. In addition, a peak centered at 2955 cm^-1^, attributed to the asymmetric stretching vibration of CH_3_ present in the RDs,^29^ was detected in all mPDC/RD composites but was absent in the pristine mPDC. Furthermore, the characteristic C=C stretching vibration at 1637 cm^-1^ and 810 cm^-1,30^ which was present RD monomers, disappeared after photopolymerization in all samples, indicating high (meth)acrylate conversion. Therefore, these FTIR results confirm the successful incorporation of IBOA, IDA, and BTA into the mPDC polymer network following photopolymerization.

**Figure 2.**
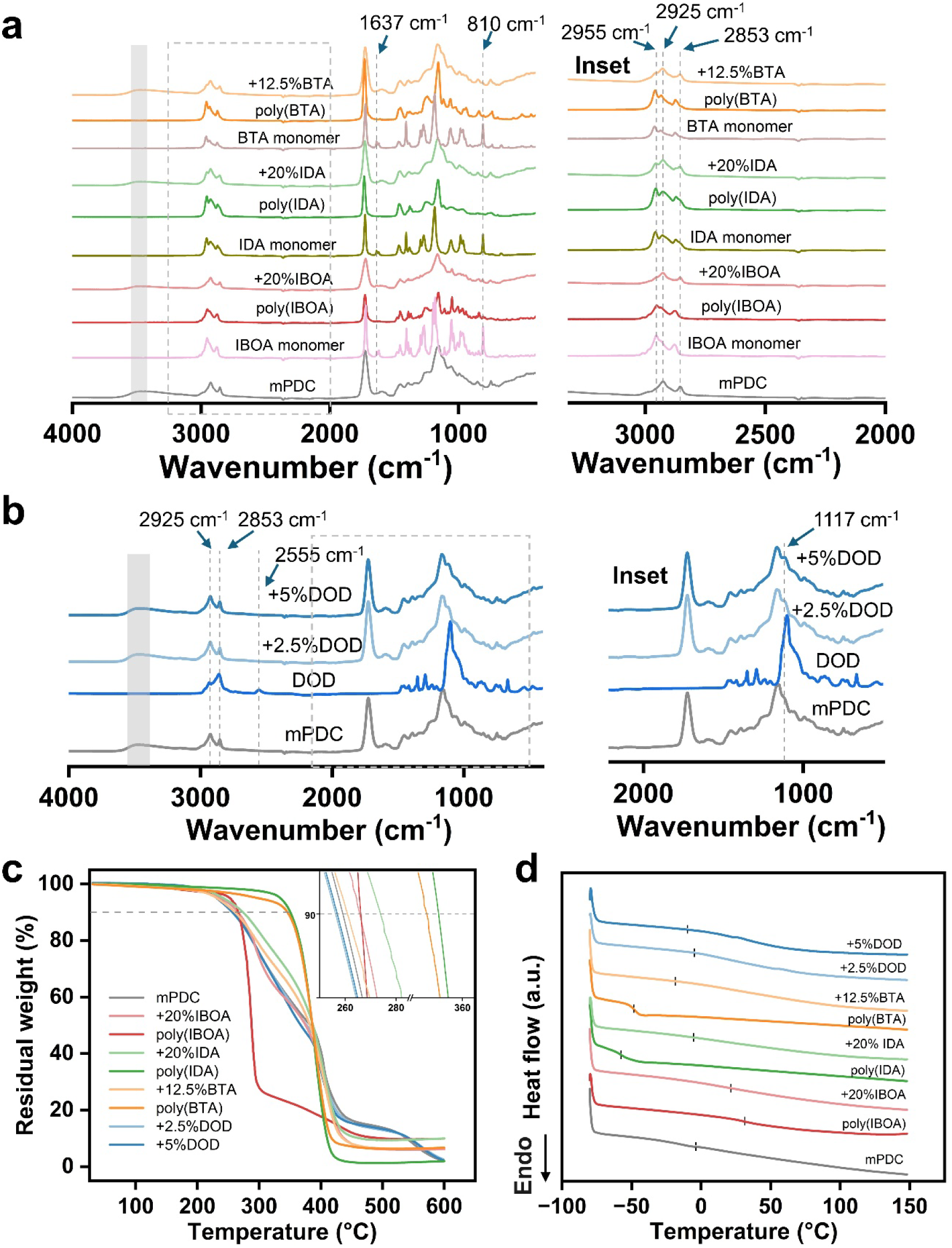
Characterization of the chemical structure and thermal properties of photopolymerized mPDC composites containing RDs and/or DOD. (**a**) FTIR spectra of mPDC+20%IBOA, mPDC+20%IDA, mPDC+12.5%BTA, with pristine mPDC, RD monomers and RD homopolymers as comparison (left). Enlarged FTIR spectra of all groups in the range of 2000-3200 cm^-1^ (right). (**b**) Left: FTIR spectra of mPDC+2.5%DOD and mPDC+5%DOD, with pristine mPDC as comparison. Right: Enlarged FTIR spectra of all groups in the range of 500-2200 cm^-1^. (**c**) TGA curves and (**d**) DSC thermograms of mPDC+20%IBOA, mPDC+20%IDA, mPDC+12.5%BTA, mPDC+2.5%DOD, mPDC+5%DOD, with pristine mPDC and the corresponding RD homopolymers as controls.

The mPDC/DOD composites were prepared by formulating inks containing 2.5 wt% and 5 wt% DOD (**Supplementary Information Table S1**). FTIR analysis revealed a distinct evolution in the band shape within the C–O–C stretching region ∼1117 cm⁻¹,^31^ indicating successful incorporation of ether-containing segments from DOD and an altered local chemical environment within the composite network (**Figure 2b**). In addition, the characteristic S-H stretching vibration at 2555 cm⁻¹,^32^ observed in the DOD monomer, disappeared after photopolymerization in mPDC/DOD composites, suggesting effective consumption of thiol groups during network formation. These FTIR results confirm the incorporation of DOD and its chemical participation in modulating the polymer network during photopolymerization.

The thermal properties of mPDC/RD composites were evaluated using thermogravimetric analysis (TGA) and differential scanning calorimetry (DSC). The TGA curves showed that pristine mPDC exhibited a decomposition temperature (T_d_) of 259.1°C (**Figure 2c**). Incorporation of RDs shifted the decomposition temperature to higher values, with T_d_ values of 265.7 °C, 273.9 °C, and 260.9 °C for mPDC/IBOA, mPDC/IDA, and mPDC/BTA, respectively. This increase in thermal stability is consistent with the relatively high decomposition temperatures of the corresponding RD homopolymers (**Supplementary Information Table S2**), which exhibited T_d_ values of 266.1 °C, 350.6 °C, and 346.0 °C for poly(IBOA), poly(IDA), and poly(BTA), respectively. In contrast, incorporation of DOD slightly reduced the decomposition temperature to 256.3 °C and 256.9 °C for mPDC+2.5%DOD and mPDC+5%DOD, respectively, which may be attributed to the introduction of flexible thioether linkages and a decrease in crosslinking density within the polymer network.

DSC thermograms (**Figure 2d**) revealed glass transition behavior for all polymers without discernible melting or crystallization peaks, indicating that these polymers were largely amorphous. Here, the glass transition temperature (T_g_) was determined by the inflection point of the DSC heat flow curve, which is particularly suitable for polymer networks in this study exhibiting broad transition regions. The T_g_ of photopolymerized mPDC was determined to be −3.4°C, consistent with previously reported values.^23^ In comparison, homopolymerized IBOA, IDA, and BTA, i.e. poly(IBOA), poly(IDA), and poly(BTA), exhibited T_g_ values of 31.5 °C, −57.7 °C, and −48.8 °C, respectively (**Supplementary Information Table S2**). Incorporation of IBOA elevated the polymer T_g_ to 20.9 °C, whereas incorporation of IDA and BTA reduced T_g_ to −18.9 °C and −5.6 °C, respectively. Notably, the mPDC/RD composites exhibited a substantially broadened glass transition region compared to pristine mPDC and the RD homopolymers, suggesting the formation of copolymerized networks and/or phase-mixed polymer blends.^33, 34^ Incorporation of DOD shifted T_g_ to lower values of −4.6 °C (mPDC+2.5%DOD) and −9.4 °C (mPDC+5%DOD).

### 2.3 mPDC composites exhibit tunable mechanical properties

Tunable mechanical properties are critical for biomaterials intended for diverse applications in regenerative engineering. Uniaxial tensile test was performed on fully hydrated, photopolymerized dog-bone-shaped specimens of mPDC composites. The results showed incorporation of RDs and/or DOD enabled substantial modulation of mechanical performance (**Figure 3** and **Supplementary Information Table S3**). Notably, all mPDC/RD composites exhibited simultaneous increases in stiffness and ductility, resulting in significantly improved toughness compared to pristine mPDC. Among all formulations, mPDC+20 wt% IBOA composite demonstrated the most pronounced improvement, exhibiting a 3.6-fold increase in Young’s modulus (or stiffness, 50.1 MPa vs. 14 MPa), an approximately 4-fold increase in ultimate tensile strength (UTS) (6.8 MPa vs. 1.8 MPa), and a 3-fold increase in strain at break (39% vs. 14%). Increasing the IBOA content to 40 wt% (mPDC+40%IBOA) further enhanced stiffness and strength, yielding nearly 10-fold increases in both Young’s modulus (134 MPa vs. 14 MPa) and UTS (18.3 MPa vs. 1.8 MPa), while still maintaining a ∼2-fold increase in strain at break (27% vs. 14%). Incorporation of IDA or BTA also improved mechanical performance, resulting in ∼4-fold increases in UTS and 2-3-fold increases in strain at break compared to pristine mPDC.

**Figure 3.**
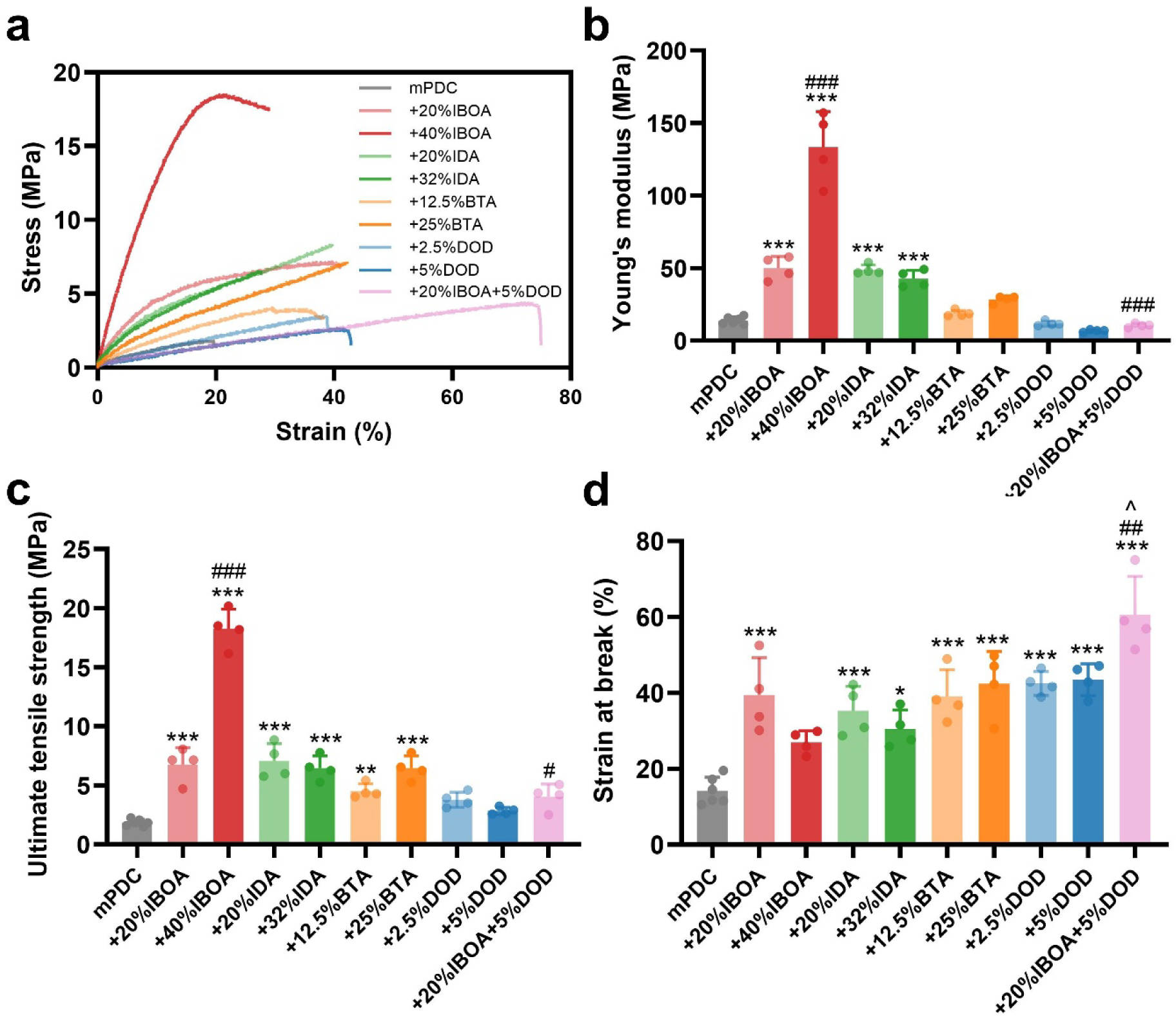
Mechanical properties of mPDC composites containing RD and/or DOD. (**a**) Representative stress-strain curves, and quantitative comparisons of (**b**) Young’s modulus, (**c**) ultimate tensile strength, and (**d**) strain at break for fully hydrated, photopolymerized composites. All samples were incubated in PBS (pH 7.4) at room temperature overnight to achieve full hydration prior to mechanical testing (n ≥ 4). Statistical significance: * vs pristine mPDC, # vs mPDC+20%IBOA, and ^ vs mPDC+5%DOD; *, # or ^ for *p* < 0.05, **, ## for *p* < 0.01, ***, ### for *p* < 0.001.

In contrast to RDs, incorporation of DOD as a chain transfer agent decreased stiffness while markedly enhancing ductility. Specifically, mPDC+5%DOD exhibited a 2-fold reduction in Young’s modulus (6.8 MPa vs. 14 MPa) but a ∼3-fold increase in strain at break (43% vs. 14%) compared to pristine mPDC. Notably, combining IBOA and DOD produced a highly ductile composite, achieving a strain at break of 61% while maintaining a Young’s modulus of 10.6 MPa, highlighting a synergistic effect arising from RD-mediated modulation of crosslinking density together with thiol-driven refinement of network topology.

Pristine photopolymerized mPDC is characterized by substantial brittleness and limited mechanical tunability. Here, incorporation of RDs and/or DOD enabled broad and continuous tuning of mechanical properties, spanning Young’s modulus from 6.8 to 134 MPa, UTS from 1.8 to 18 MPa, and strain at break from 14% to 61%. Importantly, the concurrent enhancement of stiffness and ductility observed in mPDC/RD composites suggests a toughening mechanism, rather than a conventional stiffness-ductility trade-off. The most significant toughening was observed in the mPDC/IBOA composites, which is likely attributed to several factors. First, incorporation of IBOA increased the polymer glass transition temperature from −3.4 °C to 20.9 °C (**Figure 2c**), shifting the polymer mechanical response under ambient conditions from a rubbery regime toward the glass-rubber transition region, which is often associated with an optimal balance between polymer chain mobility and restriction, and improved toughness. Second, the rigid isobornyl pendant group introduced by IBOA, together with possible formation of IBOA homopolymer-poly(IBOA) domains within the network, may restrict polymer chain mobility and enhance cohesive strength, resulting in increased stiffness and strength.^35,36^ Third, the monofunctional nature of IBOA increased average chain length between crosslinking points (**Figure 1b)**, enabling greater network extensibility and flexibility. This observation is consistent with previous reports showing that the addition of IBOA can simultaneously improve stiffness and ductility in a photopolymerizable resin.^37^ By contrast, the reduced stiffness and enhanced ductility observed upon incorporation of DOD can be attributed to its chain transfer activity. DOD is expected to shift the polymerization mechanism from purely methacrylate chain-growth toward the mixed-mode involving both methacrylate chain-growth and thiol-ene step-growth polymerization, which typically yields a more homogenous crosslinked network, thereby improving ductility.^25, 38, 39^

### 2.4 mPDC composites show diverse degradation behavior

The degradation behavior of mPDC composites was evaluated by measuring mass loss during incubation in 0.1 M NaOH aqueous solution (pH=13, 37 °C) as an accelerated degradation approach, and by assessing changes in mechanical properties after incubation in PBS (pH=7.4, 37 °C) as a physiologically relevant condition, respectively (**Figure 4**). All mPDC composites exhibited gradual mass loss at different rates, confirming their hydrolytic degradability (**Figure 4a**). Compared to pristine mPDC, incorporation of DOD, including mPDC+5%DOD and mPDC+20%IBOA+5%DOD composites, significantly accelerated degradation, which is likely attributed to increased water update and accessibility to ester bonds on the polymer chain within the thiol-acylate network. By contrast, mPDC/RD composites showed reduced degradation rates compared to pristine mPDC, suggesting that incorporation of RDs decreases water penetration and limits hydrolytic access to the ester linkages. In addition, mechanical characterization revealed no substantial changes in Young’s modulus, UTS, and strain at break for any composite before and after incubation in a PBS (pH = 7.4, 37 °C) for 28 days (**Figure 4 b-d**). The results indicated that degradation of mPDC composites proceeds slowly under physiological conditions to preserve mechanical integrity over at least four weeks. Since biomaterial scaffolds with tunable degradation profiles that match new tissue formation are highly desirable yet challenging to achieve,^40^ these findings demonstrate that incorporation of RDs and/or thiol molecules provides an effective strategy to modulate the degradation behavior of photopolymerizable CBPs.

**Figure 4.**
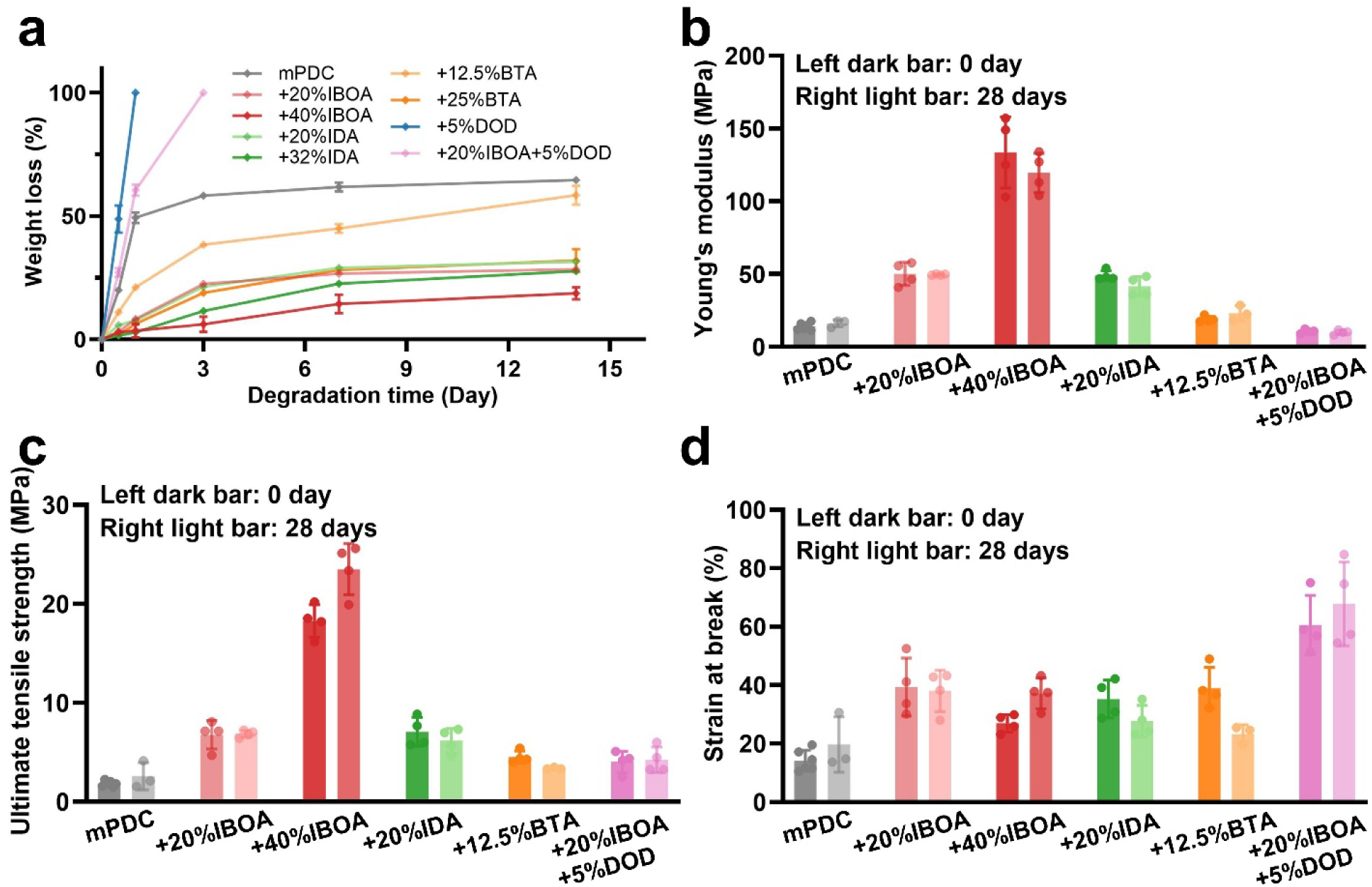
Biodegradation behavior of photopolymerized mPDC composites containing RDs and/or DOD. (**a**) Accelerated degradation profiles of mPDC-based polymers in 0.1 M NaOH aqueous solution at 37°C over 2 weeks (n=4). Mechanical characterization of (**b**) Young’s modulus, (**c**) ultimate tensile strength, and (**d**) strain at break after incubation in PBS (pH = 7.4) at 37°C for 28 days (n≥3).

### 2.5 Photopolymerized mPDC composites exhibit good cytocompatibility

Cytocompatibility is a critical prerequisite for the safe and effective use of biomaterial scaffolds. Although the excellent biocompatibility of mPDC has been previously demonstrated both *in vitro* and *in vivo*,^20, 21, 24^ it remains important to evaluate whether incorporation of RDs and/or DOD introduces any cytotoxic effects. Therefore, cytocompatibility of photopolymerized mPDC composites was assessed using both indirect extract testing and direct contact assays following the ISO 10993-5:2009 standard (**Figure 5**). Mouse fibroblast L929 cells incubated with leachable extracts from photopolymerized mPDC composites exhibited comparable cell viability and normal morphology relative to cells cultured in growth medium (**Figure 5a and 5b**). Similarly, L929 cells that were directly seeded and grown on photopolymerized mPDC composites showed no significant difference in cell viability or morphology compared to those grown on pristine mPDC (**Figure 5c and 5d**). These results indicated that incorporation of RDs and/or DOD did not compromise the cytocompatibility of mPDC, and all mPDC composite formulations displayed good cytocompatibility.

**Figure 5.**
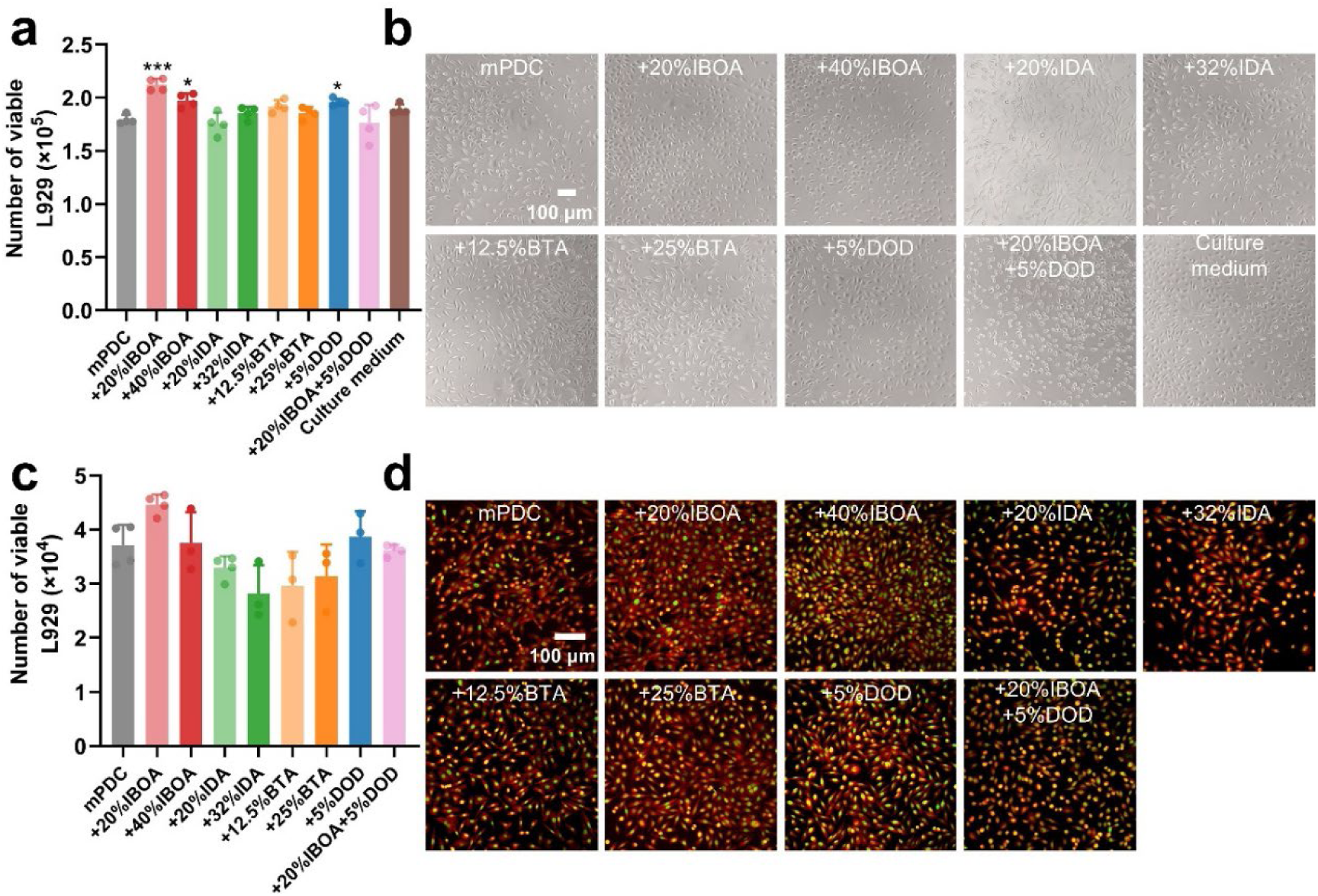
Cytocompatibility evaluation of mPDC composites. (**a**) Number of viable L929 cells quantified by AlamarBlue assay after incubation with polymer extracts (n=4). * indicates statistical significance compared to pristine mPDC group. (**b**) Representative morphologies of L929 cells cultured in extracts from various mPDC composites for 24 h. (**c**) Number of viable L929 cells (n=4) and (**d**) representative morphologies of L929 cells directly seeded and grown on surfaces of mPDC composites, visualized by F-actin/nuclei staining.

### 2.6 3D printing of meniscus scaffolds using mPDC composite inks and their biological functionality

The limited intrinsic healing capability of the meniscus, combined with its complex loading environment and exquisite anisotropic structure, makes meniscus repair or regeneration extremely challenging.^41^ 3D printing of scaffolds with biomimetic mechanical properties and structural features provides a promising strategy to restore mechanical function and promote fibrocartilage regeneration.^42–44^ To demonstrate the feasibility of mPDC composite inks for meniscus scaffold fabrication, we evaluated the 3D printability and biological functionality of selected formulations, including pristine mPDC, mPDC+20%IBOA, mPDC+20%IBOA+5%DOD, and mPDC+20%IDA (**Supplementary Information Table S1**). These formulations were selected because their tensile properties are comparable to those of native meniscus tissue.^41, 45^ Meniscus scaffolds (front/rear chord lengths: 3.45mm and 6.22 mm; width: 5.33 mm; and height: 2.52 mm) with radially and circumferentially aligned fibers (diameter: 0.36 mm; spacing: 0.36 mm) were successfully fabricated using all selected composite inks and with a home-built CLIP-based 3D printing system (**Figure 6a and Supplementary Information Figure S2a**). Notably, composite inks containing IBOA (mPDC+20%IBOA, mPDC+20%IBOA+5%DOD) required lower UV dosage to achieve optimal printing fidelity, i.e. producing scaffolds closely matching the CAD model dimensions, compared to pristine mPDC and mPDC+20%IDA inks. This is likely attributed to the higher photopolymerization reactivity of IBOA.

**Figure 6.**
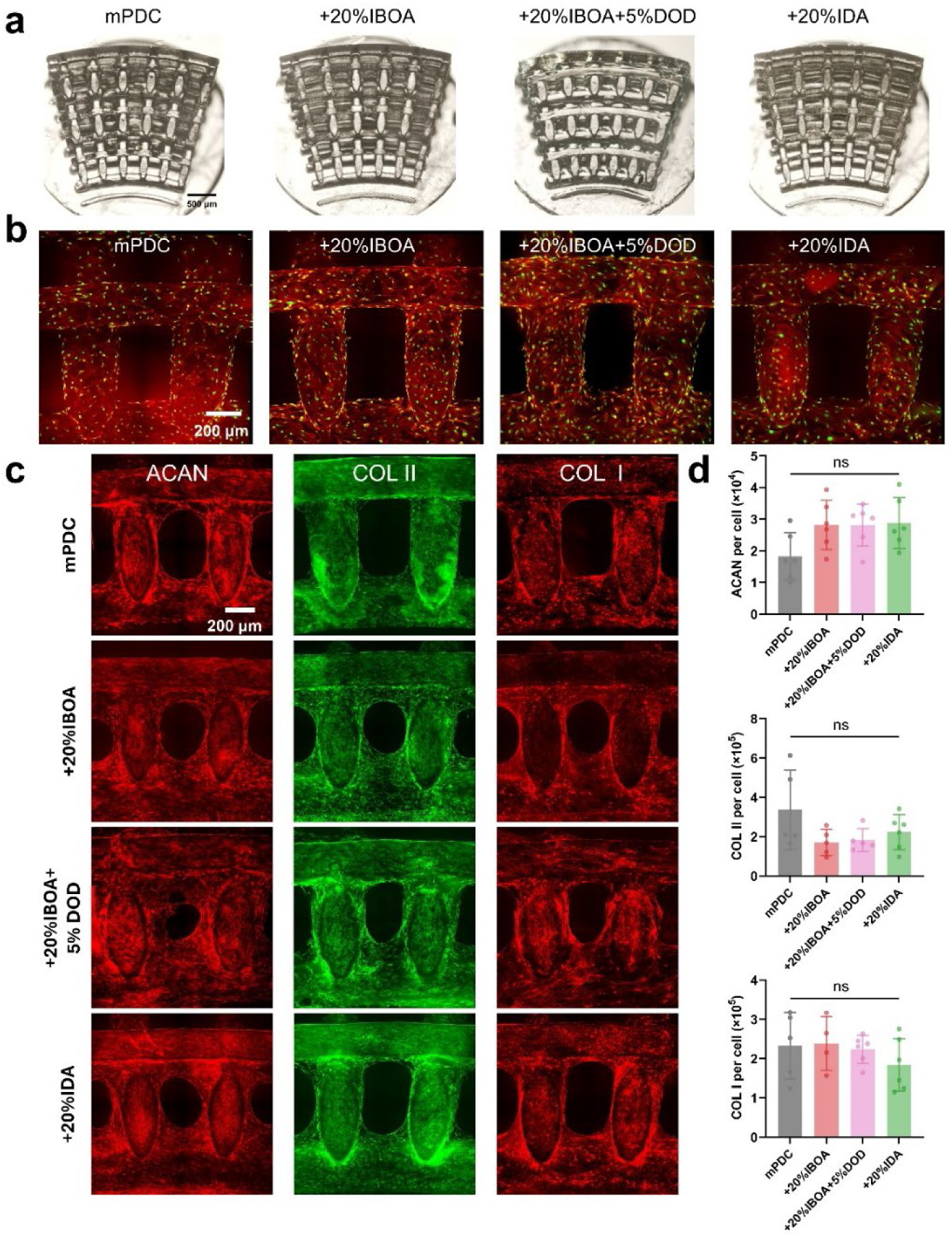
3D-printed meniscus scaffolds fabricated using mPDC composite inks support hACs growth and robust fibrochondrogenic matrix deposition. (**a**) Representative optical microscope images of 3D-printed meniscus scaffolds. (**b**) Representative immunofluorescent images (Red: F-actin; Green: Nuclei) of hACs grown on 3D-printed scaffolds. (**c**) Representative immunofluorescent images and (**d**) quantitative analysis showing fibrochondrogenic extracellular matrix deposition by hACs on 3D-printed scaffolds, including aggrecan (ACAN), collagen type II (COL II), and collagen type I (COL I).

To assess their biological functionality, human articular chondrocytes (hACs) were seeded and grown within the 3D-prined meniscus scaffolds. Fluorescence imaging revealed that hACs exhibited normal attachment, spreading, and growth on all mPDC composite scaffolds (**Figure 6b**), consistent with the high cell viability observed across all groups (**Supplementary Information Figure S3**). Interestingly, scaffolds fabricated from mPDC+20%IBOA and mPDC+20%IDA supported slightly higher cell viability compared to pristine mPDC and mPDC+20%IBOA+5%DOD. Following culture in fibrochondrogenic differentiation medium supplemented with 10 ng/mL transforming growth factor β3 (TGF-β3) for 7 days, hACs within all scaffold groups exhibited robust extracellular matrix deposition, including aggrecan (ACAN), collagen type II (COL II), and collagen type I (COL I) (**Figure 6c, 6d**). This expression profile is consistent with the fibrochondrocyte-like phenotype characteristic of native meniscus.^46–48^ No significant differences in matrix deposition were observed among different scaffold compositions, indicating that incorporation of RDs and/or DOD did not compromise the biofunctionality of pristine mPDC scaffolds.

Collectively, these results demonstrate that 3D-printed mPDC composite scaffolds provide a supportive microenvironment for chondrocyte growth and fibrochondrogenic differentiation, highlighting their strong potential for meniscus tissue engineering. Further studies are warranted to evaluate whether these scaffolds can recapitulate the complex mechanical behavior and long-term biological functionality of native meniscus during polymer degradation.

### 2.7 3D printing of vascular stents using mPDC composite inks and their biological functionality

Bioresorbable vascular stents offer a promising alternative to permanent metallic stents for the treatment of coronary and peripheral artery disease in interventional cardiology.^49, 50^ High mechanical strength and a low profile with struct thickness below 100 µm are critical for achieving favorable long-term patency of vascular stents. To demonstrate the potential of mPDC composite inks for vascular stent fabrication, we evaluated the 3D printability and biological functionality of selected formulations, including pristine mPDC, mPDC+40%IBOA, and mPDC+20%IDA, as these composites exhibited high mechanical strength. Vascular stents (inner diameter: 2.18 mm; length: 10 mm) were successfully 3D-printed in high printing fidelity using all selected composite inks with a CLIP-based 3D printing system (**Figure 7a** and **Supplementary Information Figure S2b**). Each 10 mm-long vascular stent with a strut thickness of ∼100 µm was printed within 12 minutes, demonstrating the excellent printability of these mPDC composites, characterized by both high resolution and rapid fabrication speed.

**Figure 7.**
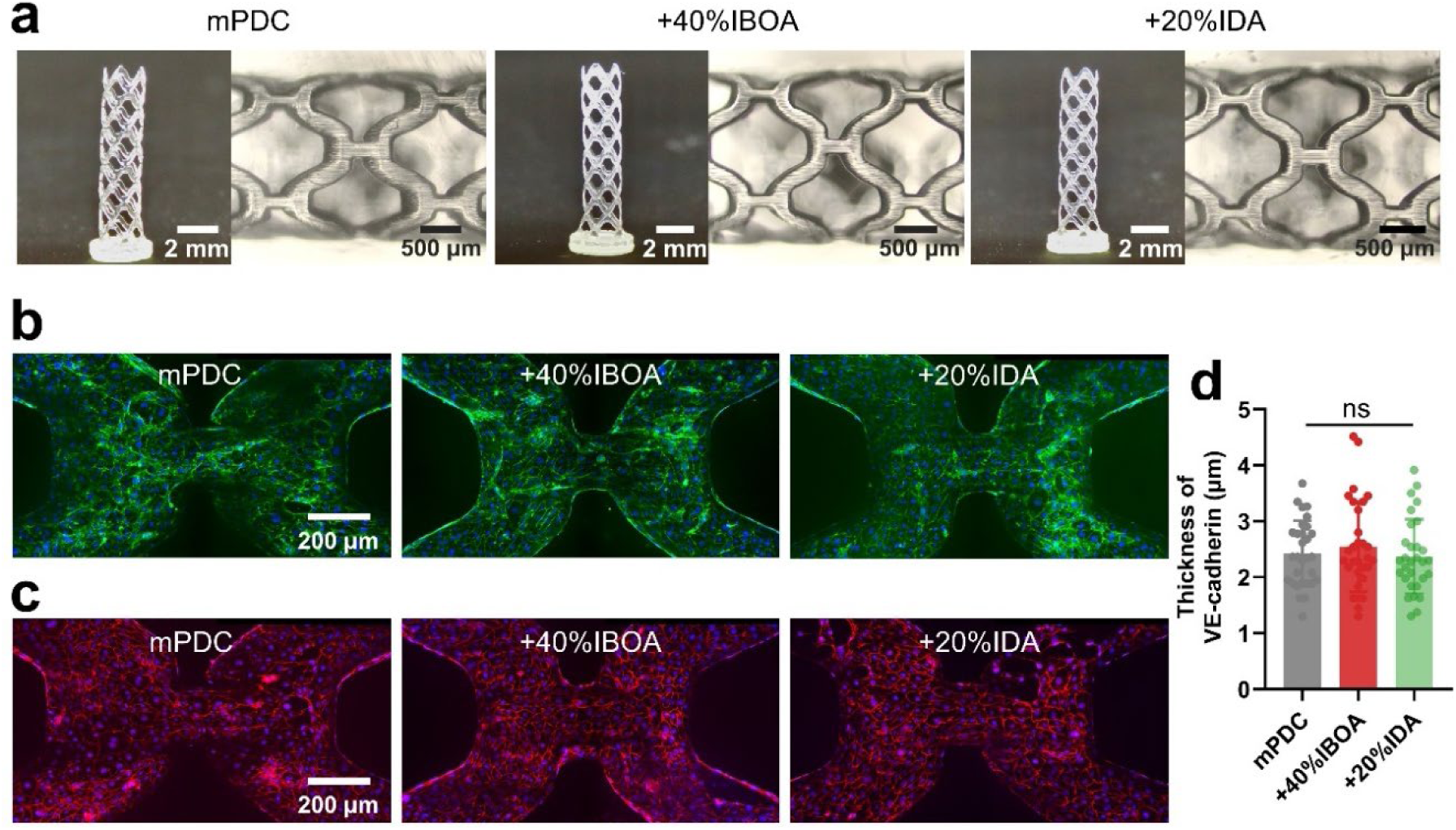
3D-printed vascular stents using mPDC composite inks support favorable interactions with endothelial cells. (**a**) Representative optical images of 3D-printed vascular stents using mPDC composite inks. (**b**) Representative immunofluorescent images (Green: F-actin; Blue: Nuclei) of HUVECs grown on 3D-printed vascular stents. (**c**) Representative immunofluorescent images of endothelial junctions (Red: VE-cadherin; Blue: Nuclei) and (**d**) quantification of endothelial junction thickness on 3D-printed vascular stents.

Rapid endothelialization, defined as recovery of an intact endothelial cell layer on the stent surface, is essential for long-term vascular stent performance. To evaluate endothelialization on mPDC composites, human umbilical vein endothelial cells (HUVECs) were seeded onto luminal surfaces of 3D-pritned vascular stents. After 3 days of culture, HUVECs showed nearly complete surface coverage and formed a continuous endothelial monolayer formation on all stent formulations (**Figure 7b**), which is consistent with the high cell viability observed across all groups (**Supplementary Information Figure S4**) Furthermore, immunostaining of VE-cadherin, a key endothelial junctional maker, revealed intact and well-organized intercellular junctions across all 3D-printed vascular stents (**Figure 7c**). Quantitative analysis of VE-cadherin staining showed comparable endothelial junction thickness among all stent groups (**Figure 7d**), consistent with values reported for healthy endothelial cells in our previous study^51^. Together, these results demonstrate that 3D-printed mPDC composite stents support favorable endothelial cell adhesion, coverage, and junction formation, highlighting their strong potential as bioresorbable vascular stents.

## 3. Conclusion

In summary, incorporation of mono-acylate RDs and/or the thiol molecule DOD into photopolymerizable CBP, such as mPDC, provides effective network-engineering strategies to modulate crosslinking density and network topology, thereby enabling highly tunable mechanical properties and degradation behavior. A series of mPDC composites containing RDs (IBOA, IDA, and BTA) and/or DOD were fabricated via photopolymerization and systematically evaluated in terms of mechanical properties, degradation behavior, cytocompatibility, 3D printability, and *in vitro* biological functionality. Incorporation of RDs and/or DOD enabled broad tunability of mechanical properties, with Young’s modulus (6.8-134 MPa), UTS (1.8-18 MPa), and strain at break (14%-61%). Notably, RD incorporation enhanced both the stiffness and ductility of mPDC composites, while DOD introduction significantly improved their ductility. All mPDC composites exhibited gradual degradation while maintaining comparable mechanical properties over 4 weeks under physiological conditions and demonstrated good cytocompatibility. Furthermore, their excellent 3D printability and potential application in regenerative engineering were demonstrated through successful 3D printing of meniscus scaffolds and vascular stents, which supported favorable cellular response of human chondrocytes and vascular endothelial cells, respectively. Collectively, these mPDC composites represent a versatile photopolymerizable biomaterial platform with broadly tunable mechanical properties, controlled degradation behavior, good cytocompatibility, and strong potential for diverse tissue repair and regeneration applications.

## Experimental sections

### 4.1 Materials

Citric acid and imidazole were obtained from Sigma-Aldrich. 1,12-Dodecanediol was purchased from Tokyo Chemical Industry (TCI). Tetrahydrofuran (THF) was supplied by Alfa Aesar and used as a solvent for polymer synthesis. Glycidyl methacrylate (GMA) was obtained from Sigma-Aldrich and used as a functional monomer. Absolute ethanol was purchased from Fisher Bioreagents and used as a solvent. For mono-functional reactive diluents (RDs), isobornyl acrylate (IBOA) was purchased from Sigma-Aldrich, and isodecyl acrylate (IDA) and butyl acrylate (BTA) were acquired from TCI. 3,6-dioxa-1,8-octanedithiol (DOD) was obtained from TCI and used as a chain transfer agent. Phenylbis(2,4,6-trimethylbenzoyl)phosphine oxide (Irgacure 819), a UV photoinitiator, was purchased from Sigma-Aldrich. Deuterated dimethyl sulfoxide (DMSO-d₆) was obtained from Sigma-Aldrich and used for nuclear magnetic resonance (NMR) spectroscopy. All chemicals were used without further purification unless otherwise specified.

### 4.2 Preparation of photopolymerizable resins

Photopolymerizable mPDC was synthesized by following a previously reported protocol.^24^ Initially, citric acid and 1, 12-dodecanediol at molar ratio of 2:1 were melted (165°C, 20 min), co-polymerized (140°C, 45 min), purified and freeze-dried to yield PDDC (poly(1, 12-dodecamethylene citrate)) prepolymer. Then, each 22 g of the PDDC prepolymer was dissolved in 180 mL of tetrahydrofuran with 816 mg of imidazole and 17.04 g of glycidyl methacrylate, heated (60°C, 6 h) and purified to yield mPDC prepolymer. The characterization of mPDC prepolymer was performed by proton nuclear magnetic resonance (^1^H NMR) using deuterated dimethyl sulfoxide (DMSO–d_6_) as the carrier solvent.

To formulate the photopolymerizable resins for subsequent polymerization, 50 wt% mPDC resins was blended with varying weight percentages of mono-functional RDs (e.g. IBOA, IDA or BTA) and/or CTA (e.g. DOD), as well as 2.2 wt% Irgacure® 819 and absolute ethanol acting separately as photo-initiator and solvent. The weight percentages of each RD were determined according to RD/mPDC double bond molar ratios of either 1:1 or 2:1. Besides, photopolymerizable resins of pristine RD with 2.2 wt% Irgacure® 819 were prepared (e.g. poly(IBOA), poly(IDA), poly(BTA)) as controls. The precise weight percentage of each component of the photopolymerizable resins were listed in **Supplementary Information Table S1**.

### 4.3 Polymerization of photopolymerizable resins under UV light

#### 4.3.1 Fabrication of molded samples

The polymerization of mPDC-based resins was performed in dog-bone-shaped polydimethylsiloxane (PDMS) molds or cylindrical silicon molds under UV exposure for 60 s (UV intensity: 365 nm; ∼7.0 mW⩽cm^-2^), covered by a thin glass slide. After molding, dog-bone-shaped and disk-shaped samples underwent thermal curing at 80°C for 24 h. These samples were named as mPDC, mPDC+20%IBOA, mPDC+40%IBOA, mPDC+20%IDA, mPDC+32%IDA, mPDC+12.5%BTA, mPDC+25%BTA, mPDC+2.5%DOD, and mPDC+5%DOD. The polymerization of pristine RD resins was performed in cylindrical silicon molds under UV exposure for 60 s and then heated for 24 h at 80°C. These samples were named as poly(IBOA), poly(IDA), poly(BTA).

#### 4.3.2 Fabrication of 3D-printed specimens

3D printing of photopolymerizable resins were performed using a homemade micro-continuous liquid interface production (MicroCLIP) printer, equipped with an oxygen-permeable Teflon AF-2400 window (Biogeneral Inc., San Diego, CA) at the base of the resin vat. A digital micromirror device (DMD; Texas Instruments Inc., Plano, TX) dynamically patterned UV light at 365 nm. Projection optics were optimized to achieve a pixel resolution of 7.1 μm × 7.1 μm at the focal plane. CAD designs were created in SolidWorks (Dassault Systèmes, Waltham, MA), and the corresponding STL files were processed using CHITUBOX 3D slicer software to generate 2D image slices at 5 μm thickness. Full-screen images were projected onto the resin vat window for photopolymerization. The base cure time was 60 s for the initial attachment of 3D-printed parts to the fabrication platform. The power intensity and exposure time for each layer in terms of different types of resins or patterns were listed in **Supplementary Information Table S2**. After printing, the samples were rinsed with ethanol to remove residual resin and thermally cured at 80°C for 24 h.

### 4.4 Characterizations of the polymerized mPDC with RDs and/or CTA

FTIR (Fourier transform IR) spectra of polymerized mPDC, homopolymerized RDs and mPDC composites were acquired in an attenuated total reflection (ATR) mode using a PerkinElmer spectrometer. FTIR spectra were recorded for the range of 400-4000 cm^−1^ with a resolution of 4 cm^-1^. TGA (thermogravimetric analysis) of polymerized mPDC, homopolymerized RDs and mPDC composites was performed starting from 30°C up to 600°C with a heating rate of 10°C⩽ min^-1^ under nitrogen flow (20 mL ⩽ min^-1^). Approximately 10 mg samples were used. The decomposition temperature (T_d_) was determined as the temperature at which 10% weight loss of each specimen occurred.^52^ DSC (differential scanning calorimetry) analysis of polymerized mPDC, homopolymerized RDs and mPDC composites was conducted with approximately 10 mg samples sealed in aluminum pans. Samples underwent an initial heating cycle from 25°C to 150°C and then remained in this state for 5 min to remove thermal history, followed by cooling to −80°C, and a second heating cycle returning to 150°C. All heating steps used a rate of 10°C⩽min⁻¹ and cooling step used a rate of 40°C min⁻¹ under nitrogen flow (20 mL⩽min^-1^). The glass transition temperature (T_g_) was measured during the second heating run.^52^

Mechanical properties of molded dog-bone-shaped samples of polymerized mPDC and its composites were evaluated using tensile tests with a 50 N load cell (CellScale Univert S). Samples were tested at a pulling rate of 1 mm⩽min^-1^, and stress-strain data were recorded. Young’s modulus was determined from the initial slope of the stress-strain curve. Tensile strength, strain at break, and Young’s modulus values were reported based on measurements from four parallel specimens.

Biodegradability tests were performed by incubating dog-bone-shaped samples in phosphate-buffered saline (PBS, pH 7.4) solution or by incubating disk-shaped specimens in 0.1 M NaOH solution, under 37 °C with shaking at 150 rpm for predetermined time intervals. Following incubation, dog-bone-shaped samples were subjected to tensile tests under conditions identical to those described in the previous section. Disk-shaped specimens were washed with DI water and vacuum-dried overnight and then measured by weight. Mass loss was calculated by comparing the initial mass (W₀) to the mass measured at each time point (Wₜ), according to equation below:

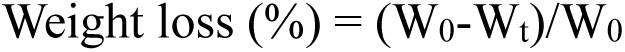

Four parallel dog-bone-shaped samples or disk-shaped samples were applied for the degradation tests.

### 4.5 Cell Culture and expansion

#### 4.5.1 Culture of mouse fibroblast cells L929

L929 (ATCC CCL-1, Manassas, VA) were cultured and expanded in growth media consisting of 10% fetal bovine serum (FBS) and 1% penicillin/streptomycin in Eagle’s Minimal Essential Medium (EMEM, Lonza, Walkersville, MD) under standard culture conditions (37°C with 5% CO_2_ in a humid environment). The media was changed every 2-3 days until the cells reached ∼80% confluency. The cells were then detached with trypsin/EDTA (Gibco/Thermo Fisher Scientific) and passaged or seeded.

#### 4.5.2 Culture of human articular chondrocytes (hACs)

Primary hACs (StemBioSys, San Antonio, TX) were cultured and expanded in flasks according to manufacturer’s instructions. They were cultured in low-glucose Dulbecco’s Modified Eagles Medium (DMEM, Gibco, Grand Island, NY, USA), supplemented with 15% FBS, 1% Glutamax, and 1% penicillin/streptomycin under the standard culture condition (37°C with 5% CO_2_ in a humid environment). The media was changed every 2-3 days until the cells reached ∼80% confluency. The cells were then detached with trypsin/EDTA (Gibco/Thermo Fisher Scientific) and passaged or seeded. For all experiments, hACs at passages 3-4 were used.

#### 4.5.3 Culture of human umbilical vein endothelial cells (HUVECs)

Primary HUVECs (ATCC, Manassas, VA) were cultured and expanded in growth media consisting of Endothelial Csd1ell Growth Kit-VEGF (ATCC PCS-100-041) in Vascular Cell Basal Medium (ATCC PCS-100-030) under the standard culture condition (37°C with 5% CO_2_ in a humid environment) to ∼80 % confluency before passaging or seeding. Media was changed every 2-3 days and HUVECs were used at passage 5-7.

### 4.6 Cell seeding, viability and morphology of cells

#### 4.6.1 Cell seeding

Different cell seeding methods were applied to disk-shaped specimens, printed meniscus scaffolds and printed vascular stents. Before seeding, all specimens were sterilized with 70% ethanol under UV exposure for 20 min and washed with DPBS three times to remove the ethanol. For seeding cells (L929 or hACs or HUVECs) on disk-shaped specimens, 25 µL of cell suspension (10.6 × 10^4^ cells/mL) were directly seeded on the surface of the disks and incubated for ∼3.5 h to allow cell attachment on disks before being transferred into a 48 well-plate with 300 µL pre-warmed growth media in each well. For seeding hACs on the printed meniscus scaffolds, the cell suspension was first mixed with an optimal concentration of GelMA/culture media solution (10 wt%) to enhance viscosity. 30 µL of the GelMA/cell suspension (100 × 10^4^ cells/mL) were directly seeded on the printed meniscus scaffolds. They were incubated for 1 hrs under room temperature and 2.5 h in the incubator to allow cell attachment on the scaffolds before being transferred into a 24 well-plate with 1 mL pre-warmed growth media in each well. For seeding HUVECs on the printed vascular stents, the vascular stents were put into a sterile silicon tube and 20 µL of HUVECs suspension (100 × 10⁴ cells/mL) were injected to fill the entire lumen of the silicon tube. After a 5-minute incubation for cell attachment, the scaffold was gently rotated by 90°, followed by a 30-minute incubation. The cell suspension was then carefully aspirated from the tube. After rotating the scaffold another 90°, another 20 µL of HUVECs suspension (100 × 10⁴ cells/ml) was injected into the lumen of the tube, and the same seeding process was repeated. Finally, the printed vascular stent entrapped in silicon tube with attached cells were transferred into pre-warmed growth media.

#### 4.6.2 Cell viability

Non-direct contact: the sterile disk-shaped specimens made of mPDC containing RDs and/or CTA were incubated with 0.5 mL of growth media for 24 h at 37°C to leach. Next, the liquid extracts of different specimens were collected and added to L929 at subconfluence in a 48-well plate for 24-hour incubation. Direct-contact: L929 or hACs or HUVECs were seeded on disk specimens and incubated for 24 h. Cells either from non-direct contact or direct contact assessments were then incubated with growth media containing Alamar blue cell viability reagent (Thermo Fisher Scientific, San Jose, CA) for 4 h. After transferring reagents into 96-well plate, the fluorescence of each well was measured at 560/590 nm (excitation/emission) using a microplate reader. The number of viable cells per well was calculated against a standard curve prepared by plating various concentrations of isolated cells, as determined by the hemocytometer.

#### 4.6.3 Cell morphology

Cells on specimens were pre-fixed with 4 % formaldehyde (Fisher Scientific, Fair Lawn) for 10 minutes and fixed for another 15 min and permeated with 0.1 % Triton X-100 for 15 min. Then, the cells were incubated with Alexa Fluor® 568 Phalloidin (10 μM, A12381, ThermoFisher Scientific) and SYTOX™ Green Nucleic Acid Stain (0.15 μM, S7020, ThermoFisher Scientific) in PBS solution with 1.5 % bovine serum albumin for 30 min. The samples were finally washed three times in PBS and mounted on microscope dish for examination using Leica spinning disk confocal microscope. Cells cultured in the well-plate were imaged using an optical microscope.

### 4.7 Chondrocyte differentiation

Chondrogenic differentiation media consisted of high-glucose DMEM, supplemented with 1% ITS^TM^ Premix, 10 ng/mL TGF-β1, 100 nM dexamethasome, 40 μg/mL L-proline, 50 μg/mL ascorbate-2-phosphate, 1% Glutamax, 1% penicillin/streptomycin and 1% gentamicin. hACs on the printed meniscus scaffolds were first cultured in growth medium for 48 h, reaching high confluency. Then, the scaffolds with cells were transferred and cultured in chondrogenic differentiation media. The differentiation media was renewed every 2-3 days, and the scaffolds were tested after chondrogenic differentiation for 7 days.

### 4.8 Immunostaining and imaging of expressed markers of primary cells

To explore the hACs differentiation, the samples were blocked in 1.5% bovine serum albumin (BSA) solution for 40 min at room temperature to block non-specific background following fixation and permeation. hACs were then incubated with primary antibody for collagen I (12.8 μg/ml, MA1-26771, ThermoFisher Scientific), collagen II (10 μg/ml, PA5-99159, ThermoFisher Scientific) and ACAN (5 μg/ml, MA3-16888, ThermoFisher Scientific) overnight at 4°C. The samples were washed three times PBS and incubated with Goat anti-mouse IgG secondary antibody (2 μg/ml, A-21235, ThermoFisher Scientific) and Donkey anti-rabbit IgG secondary antibody (2 μg/ml, A-21207, ThermoFisher Scientific) in the dark for 1 h at room temperature. SYTOX™ Green Nucleic Acid Stain were used to stain nuclei. The specimens were then washed three times with PBS and imaged using Nikon spinning disk confocal microscope.

To examine the HUVECs phenotype, cells were first incubated with 1.5 % BSA for 40 minutes to block non-specific background following fixation and permeation. HUVECs were incubated with anti-VE Cadherin (or CD144) primary antibody (3.33 μg⩽ml^-1^; ab33168; Abcam Inc.) overnight at 4 °C, then both cells were incubated with SYTOX Green Nucleic Acid Stain (0.15 μM, S7020, ThermoFisher Scientific) and Alexa Fluor 594 Donkey anti-Rabbit IgG (H+L) secondary antibody (4 μg/ml, A21207, ThermoFisher Scientific) in PBS solution with 1.5 % BSA for 60 minutes at room temperature. The samples were finally washed three times in PBS. The printed vascular stent within silicon tube was placed on microscope dish for examination using Nikon spinning disk confocal microscope.

### 4.9 Statistics and reproducibility

Unless otherwise specified, data are presented as mean ± standard deviation. Statistical significance was assessed using one-way ANOVA followed by Tukey’s multiple comparison test, with a P value < 0.05 considered statistically significant. All experiments described were independently repeated at least three times with replicates to ensure reproducibility.

## Acknowledgements

This work was supported by National Institute of Health Trailblazer Award R21EB032535 to Y. Ding and startup funding of Worcester Polytechnic Institute to Y. Ding. The authors gratefully acknowledge Dr. Longkuan Xiang at WPI Bioprocess Center for technical support in sample processing.

