## Supplementary Information for "Network Modulation Enables 3D-Printed Citrate-Based Polymer Scaffolds with Broadly Tunable Mechanical Performance for Regenerative Engineering"

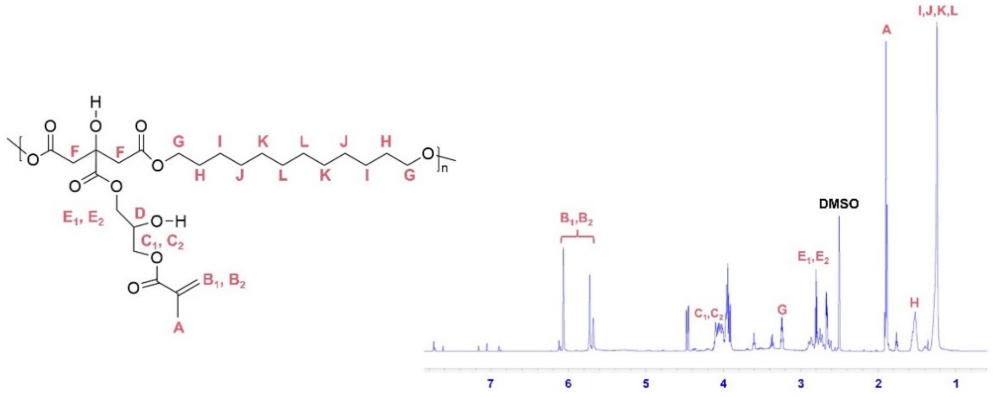


**Figure S1**. Chemical structure and NMR characterization of synthesized mPDC.

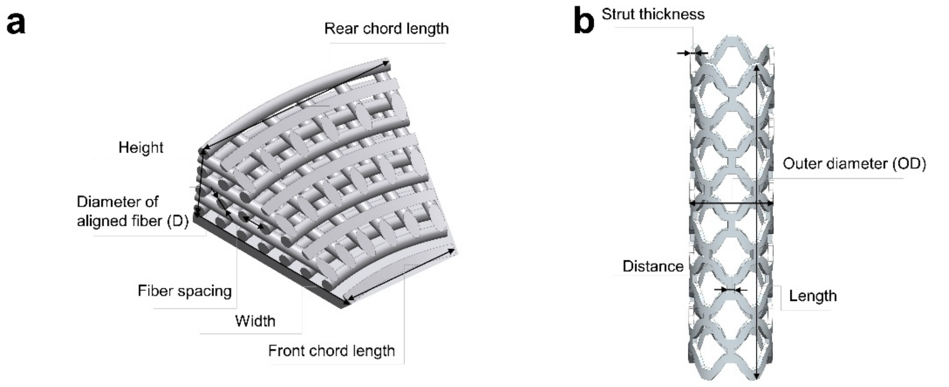


| | mPDC | +20%IBOA | +20%IDA | +20%IBOA+5%DOD | Designed meniscus model |
|---|---|---|---|---|---|
| Height (mm) | 1.28 | 1.34 | 1.39 | 1.32 | 1.40 |
| D (μm) | 201 | 217 | 181 | 225 | 200 |
| Fiber spacing (μm) | 395 | 398 | 396 | 394 | 400 |
| Width (mm) | 2.79 | 2.87 | 2.85 | 3.01 | 2.96 |
| Front chord length (mm) | 1.91 | 2.09 | 2.01 | 1.94 | 1.92 |
| Rear chord length (mm) | 3.37 | 3.63 | 3.56 | 3.38 | 3.45 |

| | mPDC | +40% IBOA | +20% IDA | Designed stent model |
|---|---|---|---|---|
| OD (mm) | 2.06 | 1.97 | 2.02 | 2.45 |
| Strut thickness (μm) | 118 | 94 | 100 | 136 |
| Distance (mm) | 0.182 | 0.163 | 0.167 | 0.230 |
| Length (mm) | 9.75 | 8.97 | 9.27 | 10.0 |

**Figure S2.** CAD models and comparison between designed and actual dimensions of (**a**) meniscus scaffold and (**b**) vascular stent.

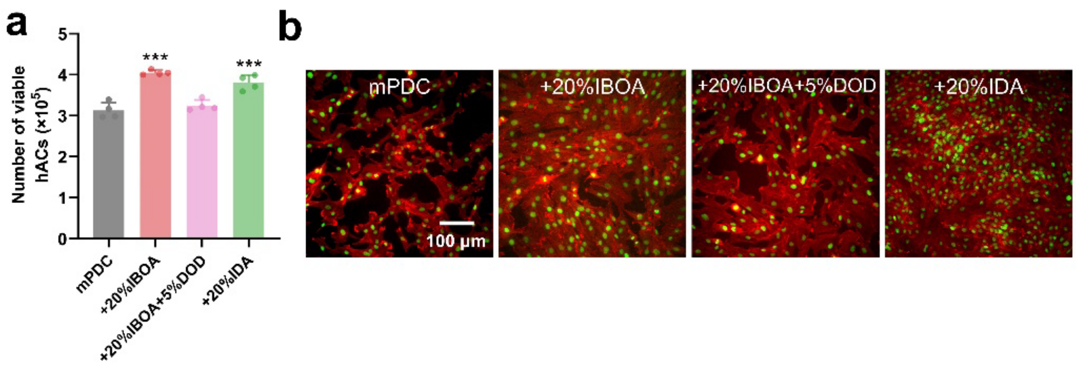


**Figure S3.** Cytocompatibility evaluation of hACs cultured on photopolymerized mPDC composites. (**a**) Number of viable cells (n=4) and (**b**) representative morphologies of hACs directly seeded and grown on mPDC composite surfaces, visualized by F-actin/Nuclei staining.

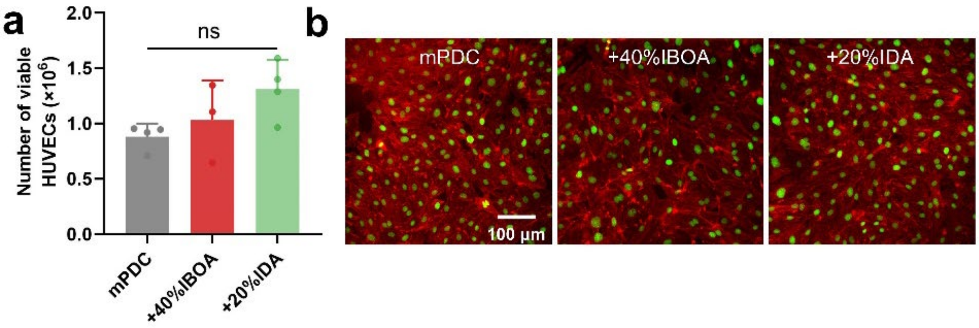


**Figure S4.** Cytocompatibility evaluation of HUVECs cultured on the photopolymerized mPDC composites. ((**a**) Number of viable cells (n=4) and (**b**) representative morphologies of HUVECs directly seeded and grown on mPDC composite surfaces, visualized by F-actin/Nuclei staining.

**Table S1.** Formulations of photopolymerizable mPDC composite inks for 3D printing.

| mPDC composite Inks | mPDC (wt%) | RDs (wt%) | DOD (wt%) | Irgacure® 819 (wt%) | Absolute ethanol (wt%) |
|---|---|---|---|---|---|
| mPDC | 50 | 0 | 0 | 2.2 | 47.8 |
| +20% IBOA | 50 | 20 | 0 | 2.2 | 27.8 |
| +40% IBOA | 50 | 40 | 0 | 2.2 | 7.8 |
| +20% IDA | 50 | 20 | 0 | 2.2 | 27.8 |
| +32% IDA[a] | 50 | 32 | 0 | 2.2 | 15.8 |
| +12.5% BTA | 50 | 12.5 | 0 | 2.2 | 35.3 |
| +25% BTA | 50 | 25 | 0 | 2.2 | 22.8 |

| | | | | | |
|---|---|---|---|---|---|
| +2.5%DOD | 50 | 0 | 2.5 | 2.2 | 45.3 |
| +5%DOD | 50 | 0 | 5 | 2.2 | 42.8 |
| +20% IBOA+5%DOD | 50 | 20 | 5 | 2.2 | 22.8 |
| pristine IBOA | 0 | 97.8 | 0 | 2.2 | 0 |
| pristine IDA | 0 | 97.8 | 0 | 2.2 | 0 |
| pristine BTA | 0 | 97.8 | 0 | 2.2 | 0 |

a: The formulation allowed for a maximum incorporation of 32 wt% IDA.

**Table S2**. Thermal properties of the photopolymerized mPDC, homopolymerized RD, mPDC+RD composites and mPDC+DOD composites.

| Thermal property | mPDC | poly(IBOA) | poly(IDA) | poly(BTA) | mPDC +20% IBOA | mPDC +20% IDA | mPDC +12.5 % BTA | mPDC +2.5% DOD | mPDC +5%D OD |
|---|---|---|---|---|---|---|---|---|---|
| $T_d$ (°C) | 259.1 | 266.1 | 350.6 | 346.0 | 265.7 | 273.9 | 260.9 | 256.3 | 256.9 |
| $T_g$ (°C) | -3.4 | 31.5 | -57.7 | -48.8 | 20.9 | -18.9 | -5.6 | -4.6 | -9.4 |

**Table S3.** Young's modulus, strain at break, and ultimate tensile strengths via tensile tests of photopolymerized mPDC composites.

| mPDC composites | Young's modulus (%) | Strain at break (%) | Ultimate tensile strength (%) |
|---|---|---|---|
| mPDC | 14.0±2.5 | 14.2±3.6 | 1.8±0.3 |
| +20% IBOA | 50.1±7.9 | 39.4±9.9 | 6.8±1.4 |
| +40% IBOA | 134±25 | 27.0±3.0 | 18.3±1.7 |
| +20% IDA | 49.0±3.4 | 35.3±6.5 | 7.1±1.5 |
| +32% IDA | 42.8±5.8 | 30.5±4.9 | 6.5±1.0 |
| +12.5% BTA | 18.9±2.1 | 39.0±7.1 | 4.5±0.6 |
| +25% BTA | 28.6±2.4 | 42.4±8.5 | 6.5±1.0 |
| +2.5%DOD | 11.9±1.7 | 42.5±3.2 | 3.8±0.6 |
| +5%DOD | 6.9±0.8 | 43.5±4.2 | 2.8±0.3 |
| +20% IBOA+5%DOD | 10.6±1.4 | 60.6±10.1 | 4.0±1.1 |

**Table S4.** 3D-printing parameters for photopolymerizable mPDC composite inks.

| mPDC composite inks | Power intensity | Exposure time (s) | Layer thickness (μm) | CAD model |
|---|---|---|---|---|
| mPDC | 1.4 | 0.62 | 5 | meniscus |
| +20% IBOA | 0.5 | 0.62 | 5 | meniscus |
| +20% IDA | 1.4 | 0.62 | 5 | meniscus |

| | | | | |
|---|---|---|---|---|
| +20% IBOA+5%DOD | 0.5 | 0.62 | 5 | meniscus |
| mPDC | 0.6 | 0.33 | 5 | stent |
| +40% IBOA | 0.313 | 0.28 | 5 | stent |
| +20% IDA | 0.5 | 0.3 | 5 | stent |
